# Structurally targeted mutagenesis identifies key residues supporting α-synuclein misfolding in multiple system atrophy

**DOI:** 10.1101/2024.07.04.602104

**Authors:** Patricia M. Reis, Sara A. M. Holec, Chimere Ezeiruaku, Matthew P. Frost, Christine K. Brown, Samantha L. Liu, Steven H. Olson, Amanda L. Woerman

**Author notes:** Correspondence: Amanda L. Woerman, PhD, Department of Department of Microbiology, Immunology, and Pathology and Prion Research Center, Colorado State University, Fort Collins, CO, USA. Authors contributed equally to the manuscript (Sara A. M. Holec, Chimere Ezeiruaku, Matthew P. Frost). Department of Surgery, Division of Abdominal Transplant Surgery, Stanford University School of Medicine, Palo Alto, CA, USA. Department of Biomedical Engineering, University of Massachusetts Amherst, Amherst, MA, USA. Department of Biochemistry and Cell Biology, Dartmouth College, Hanover, NH, USA.

## Abstract

Multiple system atrophy (MSA) and Parkinson’s disease (PD) are caused by misfolded α-synuclein spreading throughout the central nervous system. While familial PD is linked to several point mutations in α-synuclein, there are no known mutations associated with MSA. Our previous work investigating differences in α-synuclein misfolding between the two disorders showed that the familial PD mutation E46K inhibits replication of MSA prions both *in vitro* and *in vivo*, providing key evidence to support the hypothesis that α-synuclein adopts unique strains in patients. Here, to further interrogate α-synuclein misfolding, we engineered a panel of cell lines harboring both PD-linked and novel mutations designed to identify key residues that facilitate α-synuclein misfolding in MSA. These data were paired with *in silico* analyses using Maestro software to predict the effect of each mutation on the ability of α-synuclein to misfold into one of the reported MSA cryo-electron microscopy conformations. In many cases, our modeling accurately identified mutations that facilitated or inhibited MSA replication. However, Maestro was occasionally unable to predict the effect of a mutation on MSA propagation *in vitro*, demonstrating the challenge of using computational tools to investigate intrinsically disordered proteins. Finally, we used our cellular models to determine the mechanism underlying the E46K-driven inhibition of MSA replication, finding that the E46/K80 salt bridge is necessary to support α-synuclein misfolding. Overall, our studies use a structure-based approach to investigate α-synuclein misfolding, resulting in the creation of a powerful panel of cell lines that can be used to interrogate MSA strain biology.

## INTRODUCTION

Synucleinopathies are a group of neurodegenerative movement disorders, including Parkinson’s disease (PD) and multiple system atrophy (MSA), that are defined by distinct clinical and pathological features arising from the accumulation of misfolded α-synuclein proteins in the brain. While PD patients develop abnormal gait, resting tremor, muscle rigidity, and REM sleep behavior disorder,^1^ MSA patients are distinguished by the presence of autonomic failure, which can present as orthostatic hypotension, difficulty controlling urinary movement, and impaired body temperature regulation.^2^ Additionally, while PD is more prevalent and patients typically develop disease in their 60s or later, MSA is a rare disease with an earlier age of onset.^1, 2^ Along with these clinical differences, PD is defined by α-synuclein accumulating into Lewy neurites or Lewy bodies (LB) within neurons,^3^ whereas the pathological hallmarks of MSA include glial cytoplasmic inclusions (GCI) in oligodendrocytes and, to a lesser extent, neuronal cytoplasmic inclusions in neurons.^4^ Interestingly, while both diseases are defined by the presence of α-synuclein inclusions,^5, 6^ mutations in the α-synuclein gene, *SNCA*, have only been identified in PD patients,^7^ providing strong support for the hypothesis that α-synuclein misfolds into at least two distinct conformations in PD and MSA patients.

Recent advances in cryo-electron microscopy (cryo-EM) enabled the resolution of misfolded α-synuclein fibrils from human patient samples, resulting in confirmation that α-synuclein adopts two distinct structures in MSA^7^ and PD^8^ patients (Fig. 1A &B). The α-synuclein fibrils isolated from MSA patient samples are defined by the presence of two heterodimeric protofilaments, both of which adopt a Greek key motif that is stabilized by a salt bridge between residues E46 and K80. Additionally, several positively charged residues project into a pocket at the protofilament interface that contains a negatively charged non-protein density.^7^ It should also be noted that three fibril structures were resolved from MSA patient samples (PDB IDs 6XYO, 6XYP, and 6XYQ), which contain variability in subunit A but a mostly consistent structure in subunit B. In contrast, the α-synuclein Lewy fold contains a single filament with a modified Greek key motif stabilized by a salt bridge between E35 and K80. The Lewy fold also contains a negatively charged non-protein density, however, in this structure it acts to stabilize the positive charges from four internal facing lysines, none of which include residue E46.^8^

**Fig. 1.**
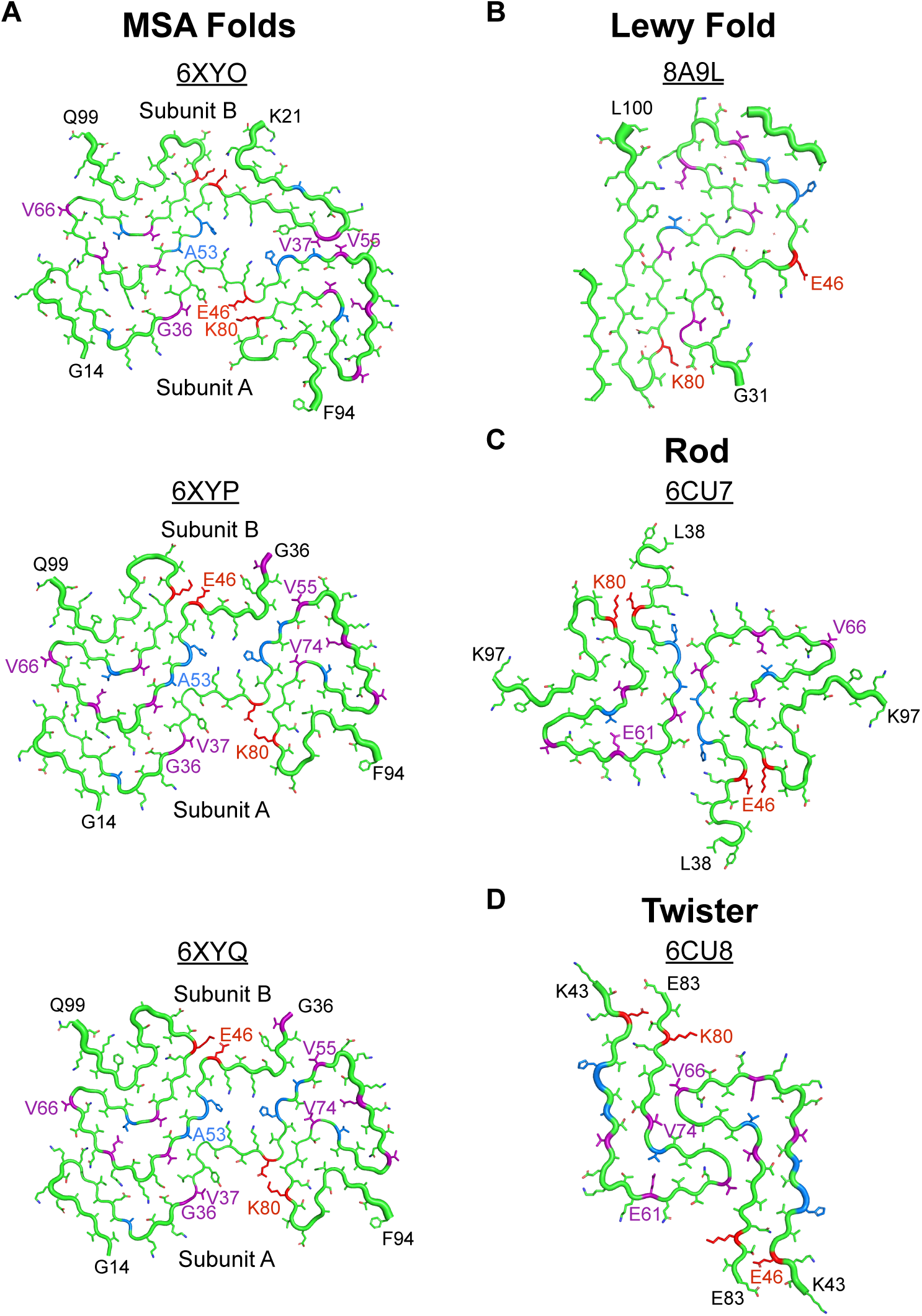
Cryo-EM structures of patient-derived and synthetic α-synuclein fibrils. The cryo-EM structures of α-synuclein fibrils resolved from (A) MSA^7^ and (B) Parkinson’s patient samples,^8^ as well as recombinant fibrils in the (C) ‘rod’ and (D) ‘twister’ conformations^25–27^ are shown. Residues that can be mutated in familial PD cases are shown in blue, with the exception of E46. Residues targeted with novel mutations designed to interfere with α-synuclein misfolding are shown in purple. The E46 and K80 residues, which often form a salt bridge that stabilizes a Greek key motif, are shown in red. PDB IDs are noted above each structure.

Using the MSA cryo-EM structures as an opportunity to develop hypotheses about how α-synuclein misfolds in disease, here we sought to test the effect of both PD-causing and novel mutations designed to interfere with protein misfolding on MSA propagation *in vitro* (highlighted in blue and purple in Fig. 1, respectively). With the goal of identifying computational tools capable of predicting the effect of mutagenesis on α-synuclein misfolding in MSA, we used Maestro software to make *in silico* predictions about the effects of each mutation on the 6XYQ MSA α-synuclein fibril structure. While total energy changes calculated using Maestro were typically capable of predicting *in vitro* outcomes, occasional discrepancies between our computational and cellular data demonstrate the challenge of using modeling software to investigate intrinsically disordered proteins that adopt a prion conformation. Additionally, varied effects of some mutations, such as the V55Y and V66F mutations, on MSA prion replication provide initial evidence that previously unreported α-synuclein conformations are likely present in a subset of MSA patients. Finally, to determine the mechanism underlying the inhibitory effect of the E46K mutation on MSA propagation, we showed that disrupting the E46/K80 salt bridge at either residue blocks cellular infection (shown in red in Fig. 1), indicating that this salt bridge is necessary to stabilize MSA prions. Unexpectedly, we could not recover MSA propagation in cells expressing the double E46K,K80E mutation, which is likely due to an increase in total energy that occurs when both residues are mutated. Overall, our studies use a structure-based approach to interrogate α-synuclein misfolding in MSA, resulting in the generation of a robust panel of cellular models that can be used more broadly to investigate α-synuclein strain biology.

## MATERIALS AND METHODS

### Human tissue samples

The Massachusetts Alzheimer’s Disease and Research Center (MADRC) supplied frozen midbrain tissues from two patients with neuropathologically confirmed MSA and two control samples. The NIH NeuroBioBank supplied tissue from two control patients (pons), and the Sydney Brain Bank supplied frozen pons samples from three neuropathologically confirmed MSA cases. Demographic information about the patient samples used is included in Table S1.

### Cell line development

Wild-type (WT) and mutant (E46K, H50Q, G51D, A53E, G36K, V37F, V55Y, E61Q, V66F, V74I, and V74P single mutations and the A30P, A53T double mutation) human α-synuclein cDNA sequences fused to enhanced yellow fluorescent protein by an 18 amino acid flexible linker (EFCSRRYRGPGIHRSPTA) were synthesized and cloned into the pcDNA3.1(+) expression vector by GenScript. The α-syn140-YFP sequences were then subcloned into the pIRESpuro3 vector (Takara) using restriction cloning with EcoRV (5’) and NotI (3’). Gene sequence and insertion were confirmed by Sanger sequencing before subsequent use. Other mutant constructs (A30G, A53V, T72M, K80E, K80N, K80Q, K80W, and the double mutant E46K,K80E) were generated by PCR amplification of the pIRESpuro3 vector containing the WT α-syn140-YFP sequence using primers designed to introduce each mutation. Gene sequence and insertion were confirmed by Sanger sequencing before subsequent use.

HEK293T cells (ATCC) were cultured in Dulbecco’s modified Eagle’s medium (DMEM; Corning) supplemented with 10% fetal bovine serum (FBS), 100 μg/mL penicillin, and 100 μg/mL streptomycin (ThermoFisher), referred to hereafter as complete media. Cultures were maintained in a humidified atmosphere of 5% CO_2_ at 37 °C. Cells were plated at a density of 5.7 × 10^5^ cells per well in a 6-well plate overnight in complete media before adding 1 μg of plasmid DNA and incubating with 3.5 μL Lipofectamine 2000 (ThermoFisher) for 20 min. Stable cells were selected in complete media containing 1 μg/mL puromycin (ThermoFisher) for 48 h before generating monoclonal lines by limiting dilution of polyclonal cells in 384-well plates. The resulting monoclonal lines were frozen in liquid nitrogen. Lysates from the lines were collected in 1× radioimmunoprecipitation assay (RIPA) buffer containing 50 mM Tris-HCl, pH 7.5 (ThermoFisher), 150 mM NaCl (Sigma), 5 mM EDTA (ThermoFisher), 1% nonidet P-40 (ThermoFisher), 0.5% deoxycholate (ThermoFisher), and 0.1% sodium dodecyl sulfate (SDS; ThermoFisher). Cell lysates in RIPA buffer were frozen, thawed, and clarified using two low-speed spins (500 × *g* for 5 min followed by 1,000 × *g* for 5 min). Total protein was measured in the supernatants via bicinchoninic acid (BCA) assay (Pierce). To compare the expression of α-syn140-YFP across the clones, a total of 10 μg total protein was prepared in 1× NuPAGE LDS loading buffer and boiled for 10 min. Samples were loaded onto a 10% Bis-Tris gel (ThermoFisher) and SDS-PAGE was performed using the MES buffer system. Protein was transferred to a polyvinylidene fluoride (PVDF) membrane. The membrane was blocked in blocking buffer [5% (wt/vol) nonfat milk in 1× Tris-buffered saline containing 0.05% (vol/vol) Tween 20 (TBST)] for 30 min at room temperature. Blots were incubated with primary antibody for GFP (1:10,000; Abcam) in block buffer overnight at 4 °C. Membranes were washed three times with 1× TBST before incubating with horseradish peroxidase-conjugated goat anti-rabbit secondary antibody (1:10,000; Abcam) diluted in blocking buffer for 1 h at 4 °C. After washing blots three times in 1× TBST, membranes were developed using the enhanced chemiluminescent detection system (Pierce) for exposure to X-ray film.

### α-Synuclein prion bioassay

Frozen brain samples were used to make a 10% (wt/vol) homogenate in calcium- and magnesium-free 1× Dulbecco’s phosphate buffered saline (DPBS) using an Omni Tissue Homogenizer with disposable plastic soft tissue tips (Omni International). Aggregated protein was isolated from the samples using phosphotungstic acid (PTA; Sigma-Aldrich) as described.^9, 10^ Briefly, the brain homogenates were incubated with 2% (vol/vol) sarkosyl (Sigma) and 0.5% (vol/vol) benzonase (Sigma) at 37 °C under constant agitation (850 RPM). Sodium PTA, dissolved in 10% (wt/vol) in double-distilled H_2_O and brought to pH 7.0, was added to a final concentration of 2% (vol/vol) and incubated overnight as described. Samples were centrifuged at 13,200 x *g* for 30 min, and the supernatants were discarded. The pellets were resuspended in 2% sarkosyl and 10% PTA before incubating for 1 h under agitation as before. Samples were centrifuged at 13,200 x *g* for 30 min, and the supernatants were removed immediately after. The pellets were resuspended in 1× DPBS (Corning) using 10% of the starting volume and were stored at 4 °C until use.

The α-syn140-YFP cell assay was performed as previously described.^10, 11^ Briefly, monoclonal subclones stably expressing WT or mutant α-synuclein were cultured in complete media in a humid 5% CO_2_ environment at 37 °C. Cells were grown in 384-well plates with black polystyrene walls (Greiner) with growth medium containing 0.012 µg/well of Hoechst 33342 (Thermo Fisher) to visualize nuclei. Plates with cells were incubated at 37 °C for 2–4 h. PTA-precipitated patient samples were diluted in 1× DPBS prior to incubation with Lipofectamine 2000 (Thermo Fisher) at room temperature for 1.5 h. Warmed OptiMEM (Thermo Fisher) was added to each sample prior to plating the sample in 6 replicate wells. Optimized assay conditions are reported in Table S2. Plates were covered with a membrane to prevent drying during incubation for 4 d at 37 °C in a humidified atmosphere with 5% CO_2_. The plate was imaged using the MicroXLS (Molecular Devices) collecting a DAPI and FITC image (for Hoechst and YFP, respectively) from 5 areas per well. The images were analyzed using MetaXpress software using parameters developed to detect and quantify intracellular aggregates in live cells using pixel intensity and size thresholds. The measure of aggregation was calculated as the total fluorescence measured across all aggregates per cell in each well (× 10^6^ arbitrary units [A.U.]).

### Modeling α-synuclein mutations on the MSA cryo-EM structure

The MSA α-synuclein filament PDB file (6XYQ) was increased to a hetero 12-mer using MOE (Chemical Computing Group) using the build_fibril.svl. From this file, Maestro (Schrödinger) was used to optimize each amino acid side using the Predict Side Chain Panel, then minimized using the OPLS4 force field. Minimization was performed to an RMSD of 10^-9^ kcal/mol/Å for convergence. From the minimized structure, each mutation was simulated manually and mutations in each heterodimeric sequence were performed independently. To maintain the same ground state energy of the protein complex, the first step in the analysis pipeline was to create a compensating mutation at an amino acid with few neighboring side chains to serve as a control (shown in Table S3). Using the V55Y mutation as an example for the analysis procedure, the minimized structure underwent a computational E83Y mutation on all 6 layers of subunit A in the total fibril structure. Using this control, we computationally made a V55Y mutation and a concomitant Y83V mutation on all 6 layers. After re-predicting the position of all side chains in proximity to both the Y55 and V83 residues using Predict Side Chain panel, both the mutated and control (E83Y) structures were minimized to convergence using the OPLS4 force field. To calculate the relative energy change for each mutation (in kcal/mol), the total calculated ensemble energies for the mutated subunits were subtracted from the structures with only the compensating mutations reported in Table S3. For V55Y, the subunit B mutations followed an identical process except that the control remained E83Y on subunit A.

### Statistical analysis

Data are presented as mean ± standard deviation. Cell assay data were analyzed using GraphPad Prism software. Mean fluorescence intensity values and standard deviations were calculated for each cell line by averaging the fluorescence intensity values from the 6 technical replicate wells. Comparisons between control and MSA patient samples were performed using the average value for each sample via Welch’s two-tailed *t*-tests with unequal variance. Significance was determined as having a *P* value <0.05.

## RESULTS

### Parkinson’s mutations exert differential effects on MSA prion replication *in vitro*

Building on our previous findings that the familial PD mutation E46K^12^ prevents MSA prion replication both *in vitro* and *in vivo,*^11, 13^ we sought to determine the effect of other known PD mutations on the replicability of MSA prions using a larger panel of α-syn-YFP cell lines. In this assay, HEK293T cells expressing α-synuclein fused to YFP via a flexible linker are incubated with α-synuclein prions isolated from deceased human patient samples. We compared 4 control samples from patients lacking neuropathological inclusions (C1, C2, C6, and C7) with 5 neuropathologically-confirmed MSA patient samples (MSA2, MSA3, MSA6, MSA7, and MSA12; Table S1) to measure α-synuclein prion aggregation in the α-syn-YFP cells (reported as fluorescence/cell × 10^6^ A.U.). Consistent with our previous results, MSA prions induced aggregation in cells expressing WT α-synuclein (*P* < 0.05) and the A30P,A53T double mutation (*P* < 0.05), but replication was inhibited in cells expressing the E46K mutation (Fig. 2A-C; *P* < 0.05).^11^ While a statistically significant difference was detected between the control and MSA samples in the E46K cells (Fig. 2B), the minimal effect size indicates the lack of a biologically meaningful result.

**Fig. 2.**
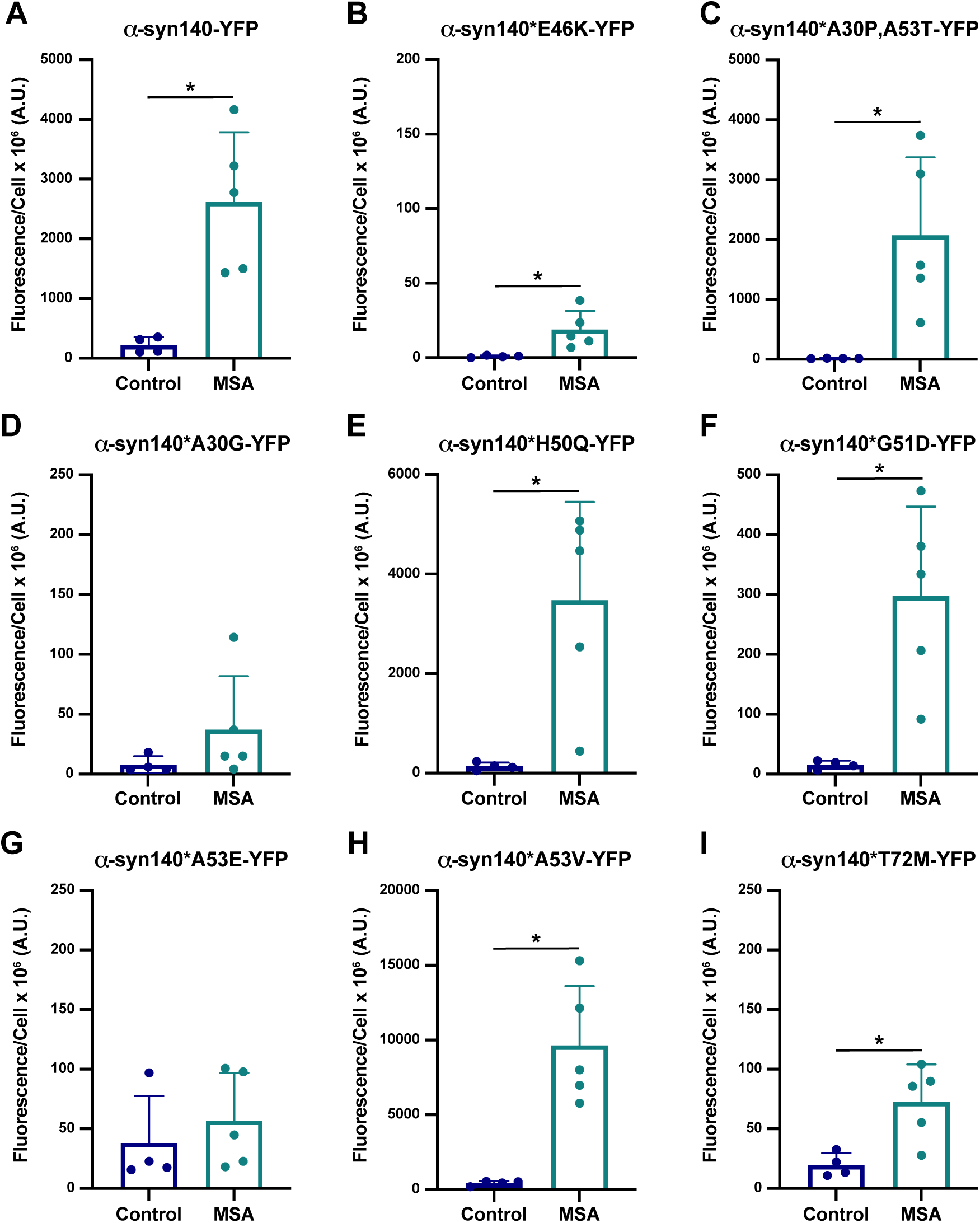
MSA prions differentially replicate in cells expressing PD-causing α-synuclein mutations. HEK293T cells expressing (A) WT α-synuclein or (B-I) Parkinson’s-causing α-synuclein mutations fused to YFP were used to determine the effect of each mutation on MSA prion replication *in vitro*. Each cell line was infected with α-synuclein prions isolated from 4 control patient samples (C1, C2, C6, and C7) or 5 MSA patient samples (MSA2, MSA3, MSA6, MSA7, and MSA12) by phosphotungstic acid precipitation (× 10^6^ arbitrary units [A.U.]). (A-C) Consistent with previous findings, (A & C) MSA samples replicated in cells expressing full-length WT α-synuclein (α-syn140-YFP) and the A30P and A53T double mutation (α-syn140*A30P,A53T-YFP), (B) whereas the E46K mutation (α-syn140*E46K-YFP) blocked MSA propagation. (D-I) More recently identified PD-causing mutations were tested to determine the effect of each on MSA propagation. The (E) H50Q (α-syn140*H50Q-YFP), (F) G51D (α-syn140*G51D-YFP), and (H) A53V mutations (α-syn140*A53V-YFP) facilitate MSA propagation, whereas the (D) A30G (α-syn140*A30G-YFP) and (G) A53E mutations (α-syn140*A53E-YFP) ablate replication. (I) The T72M mutation (α-syn140*T72M-YFP) blunted MSA prion replication compared to the α-syn140-YFP cell line. (* = *P <* 0.05)

Our initial studies focused on the first three *SNCA* mutations identified in Parkinson’s cohorts, however, several additional PD-linked mutations have since been reported. These additional mutations include A30G,^14^ H50Q,^15^ G51D,^16^ A53E,^17^ A53V,^18^ and T72M.^19^ To assess the effect of each of these PD-linked mutations on the ability of α-synuclein isolated from MSA patient samples to replicate, we first used Maestro software to measure the relative energy changes (in kcal/mol) of each mutation on the 6XYQ MSA fibril structure (Table 1) resolved via cryo-EM.^7^ Energy values below 0 after mutagenesis indicate a favorable energy change, whereas those greater than 0 indicate an unfavorable energy change. It is important to note that residue 30 is outside of the templating region of subunit B in the 6XYQ MSA fibril conformation, limiting our ability to use Maestro to evaluate the effect of the A30G mutation on MSA. However, a slight increase in total energy in subunit A suggested that the A30G mutation would reduce or inhibit MSA replication. In contrast, Maestro predicted the H50Q mutation would decrease the total energy of the MSA structure, increasing the likelihood of replication. Similar predictions were identified for the G51D mutation, though these effects were limited to subunit B as there was essentially no change for subunit A. This latter effect is intriguing given the location of residue 51 at the protofilament interface of the MSA structures. Similarly, residue 53 sits within the protofilament interface, raising the potential that mutations could disrupt replication. Here, mutagenesis of the alanine to either a glutamic acid or a valine resulted in an unfavorable increase in energy in subunit A. Notably, this increase was substantially larger for the A53E mutation compared to A53V. By comparison, the A53E mutation increased total energy for subunit B whereas the A53V mutation yielded a decrease in total energy. It is important to note that the negatively charged non-protein density that fills that pocket between the two protofilament subunits in the MSA cryo-EM structures is absent in the PDB structures. As a result, our initial *in silico* analysis of the A53E mutation resulted in a salt bridge forming between A53E and K43. *In vitro*, the presence of that density would prevent this interaction from occurring. To prevent this effect in our modeling data, we inserted a K43A mutation into the control and mutant calculations. Finally, the T72M mutation resulted in a slight increase in total energy, with a greater effect on subunit A than subunit B.

**Table 1.** Total energy change after mutagenesis using PD-causing mutations in MSA structure 6XYQ.

| Cell Line | Total Energy Subunit A | Total Energy Subunit B |
| --- | --- | --- |
| A30G | 12.2 | n.d. |
| H50Q | -19.1 | -14.8 |
| G51D | 0.5 | -11.1 |
| A53E | 30.5 | 18.5 |
| A53V | 5.0 | -6.2 |
| T72M | 10.2 | 1.5 |

*PDB structure reported previously.<sup>7</sup> Total energy reported in kcal/mol.* *n.d., not determined. Residue A30 is not present in Subunit B of the 6YXQ structure.*

To determine how well Maestro predicted the effect of α-synuclein mutations on MSA replication *in vitro*, we constructed α-syn140-YFP cell lines expressing each of the PD-linked mutations and found that the A30G mutation largely inhibited MSA replication (Fig. 2D; Table S4; *P* = 0.22), which is consistent with the mutagenesis prediction for subunit A. By comparison, the H50Q (Fig. 2E) and G51D (Fig. 2F) mutations permitted MSA replication *in vitro* (*P* < 0.05; Table S4), consistent with the favorable predictions from the Maestro analyses. Interestingly, the mutations at A53 resulted in dichotomous effects, with the A53E mutation blocking aggregation (Fig. 2G; *P* = 0.51) while the A53V mutation permitted MSA replication (*P* < 0.05; Fig. 2H; Table S4). This is notable given that predictions for the A53V mutation varied by subunit whereas the A53E analyses showed a substantial increase in total energy for both protofilament subunits. Intriguingly, while the T72M mutation did not completely inhibit MSA propagation, we observed a blunted response compared to the α-syn140-YFP cells expressing WT protein, suggesting that the T72M mutation alters replication kinetics *in vitro* (Fig. 2I; Table S4; *P* < 0.05). This is relatively consistent with the modeling data, where the mutation caused a slight increase in total energy.

Overall, the ability of Maestro to predict the effect of PD-causing point mutations on α-synuclein misfolding in MSA was reasonably accurate. Moreover, the cellular results contribute important data to the growing literature showing that the structural differences in α-synuclein fibril conformations in PD^8^ and MSA^7^ result in unique biological properties between the two strains.^11, 20–22^ Unexpectedly, we found that the effect of mutations at two residues – A30 and A53 – are mutation-specific. For example, while the A30P, A53T, and A53V mutations enable MSA replication, the A30G and A53E mutations block this activity. While an explanation for these residue-specific effects is not clear based on the mutagenesis modeling, future studies will focus on investigating this phenomenon.

### Structurally informed mutations reveal varied biological activities between MSA patient samples

Preformed fibrils (PFFs) made from recombinant α-synuclein are widely used throughout the literature as a tool to investigate synucleinopathies, including PD and MSA. Previously, we showed that PFFs made from both WT and mutant protein exhibit unique biological and biochemical properties in cell and animal models of disease^11, 13, 23^. These findings are supported by cryo-EM structures of PFFs, which misfold into conformations that are distinct from those found in PD and MSA patient samples (reviewed in Ref^24^). To further investigate the biological consequences of these structural differences, we developed a library of α-syn-YFP cell lines expressing mutations predicted to interfere with protein misfolding in either the ‘rod’ (PDB ID: 6CU7) or ‘twister’ (PDB ID: 6CU8) conformations,^25–27^ or the MSA structures.^7^ Comparing the rod and twister polymorphs (shown in Fig. 1C & D),^25^ we chose to test the effect of mutations at residues E61 and V66 on MSA replication. Our initial assessment of these two structures suggested that the E61Q mutation would alter the stability of the pocket formed by the Greek key motif (which is present in most α-synuclein structures) with a greater disruption to the twister conformation than the rod, where the residues are more tightly packed together. Similarly, mutating the valine at position 66 to a phenylalanine (V66F) would disrupt the intermolecular zipper at the protofilament interface in the twister polymorph without affecting the rod, where V66 projects outward toward solvent. We subsequently used Maestro to predict the effect of these two mutations on the MSA fibril structure and found that the E61Q mutation enhanced the energetic favorability of the MSA fibril while the V66F mutation was neutral (Table 2). Inserting these mutations into the α-syn-YFP cells, we found that all 5 MSA patient samples tested infected cells expressing the E61Q mutation (Fig. 3A; Table S5; *P <* 0.05), whereas the V66F mutation was more selective with some patient samples showing robust replication while others were completely inhibited (*P* = 0.08; Fig. 3B; Table S5). Given that the V66F mutation exhibited a neutral effect in our mutagenesis modeling (Table 2) and that the R group projects away from the fibril core in the MSA structures,^7^ these findings suggest that previously undetected α-synuclein conformations in MSA patients may exist.

**Fig. 3.**
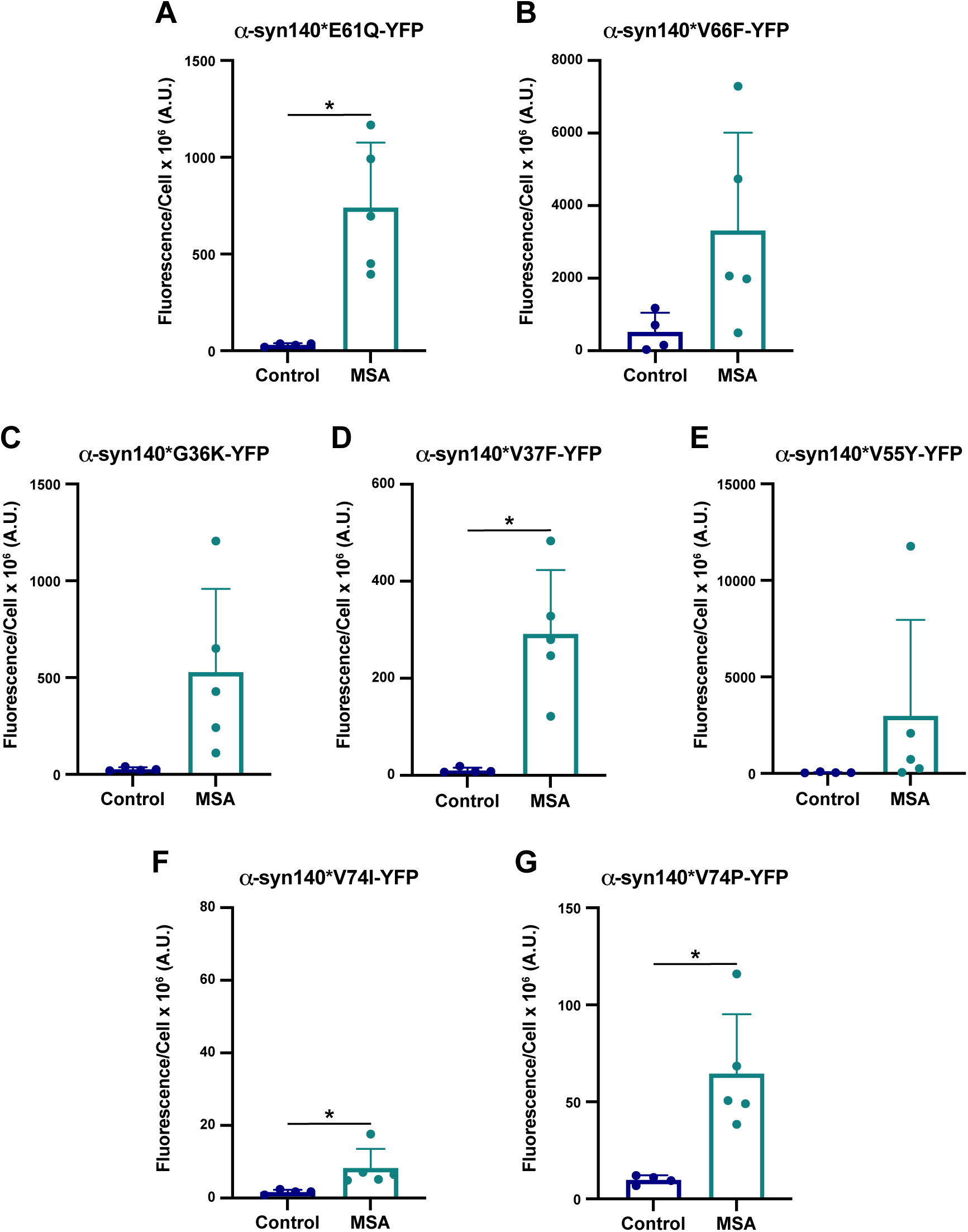
Using cryo-EM structures to inform α-synuclein mutation design reveals patient-to-patient differences in strain biology. HEK293T cells expressing α-synuclein mutations designed to interfere with protein misfolding into either the rod (PDB ID: 6CU7), twister (PDB ID: 6CU8), or MSA filaments (PDB IDs: 6XYO, 6XYP, & 6XYQ) were used to determine the effect of single residue substitutions on MSA prion replication *in vitro*. Each cell line was infected with α-synuclein prions isolated from 4 control patient samples (C1, C2, C6, and C7) or 5 MSA patient samples (MSA2, MSA3, MSA6, MSA7, and MSA12) by phosphotungstic acid precipitation (× 106 arbitrary units [A.U.]). (A) The E61Q (α-syn140*E61Q-YFP) and (B) V66F (α-syn140*V66F-YFP) mutations were designed to preferentially interfere with α-synuclein misfolding into the twister conformation. While the E61Q mutation facilitated MSA replication, successful propagation in cells expressing the V66F mutation varied by sample. (C-E) The G36K, V37F, and V55Y mutations were designed to disrupt the protofilament interface in the MSA cryo-EM structures. (C) Infection in cells expressing the G36K mutation (α-syn140*G36K-YFP) also showed patient-to-patient variability whereas the (D) V37F mutation (α-syn140*V37F-YFP) facilitated replication and (E) the V55Y mutation (α-syn140*V55Y-YFP) prevented replication of all but one MSA patient sample sample. (F & G) The V74I and V74P mutations were selected based on the hypothesis that they would interfere with the twister conformation more than the MSA structures. (F) The V74I mutation (α-syn140*V74I-YFP) blocked MSA prion replication. (G) By comparison, the V74P mutation (α-syn140*V74P-YFP) blunted MSA replication without complete inhibition. (* = *P* < 0.05)

**Table 2.** Total energy change after mutagenesis of structurally-informed mutations in MSA structure 6XYQ.

| Cell Line | Total Energy Subunit A | Total Energy Subunit B |
| --- | --- | --- |
| E61Q | -49.2 | -19.3 |
| V66F | -0.9 | 0.4 |
| G36K | 35.8 | -14.3 |
| V37F | 62.3 | -20.1 |
| V55Y | -1.5 | 35.8 |
| V74I | -13.5 | -1.7 |
| V74P | -74.7 | -76.1 |

*PDB structure reported previously.<sup>7</sup> Total energy reported in kcal/mol.*

Analyzing the three MSA fibril structures resolved by cryo-EM,^7^ we predicted that mutating residues G36, V37, and V55 would allow us to disrupt the interface between the two protofilaments in the MSA fibrils, with a particularly strong effect on 6XYO. To achieve this goal, we tested the G36K, V37F, and V55Y mutations both *in silico* and *in vitro* (Tables 2 & S4; Fig. 3C-E). Consistent with our design of these mutations, Maestro predicted the G36K and V37F mutations would cause an increase in total energy for subunit A, but a decrease for subunit B (Table 2). By comparison, the V55Y mutation was energetically favorable when we inserted the mutation into subunit A, but caused an increase in total energy for subunit B. In predictions like these where total energy between the two subunits moves in opposite directions, we predict that the disfavored conformation will exert a stronger effect than the favorable subunit, resulting in an inhibitory effect on protein misfolding. In line with our findings for the V66F mutation, the G36K mutation inhibited or slowed replication of some MSA patient samples while facilitating replication of others (*P* = 0.06; Fig. 3C; Table S5). The V37F mutation permitted replication of all MSA samples tested, though the magnitude of change was reduced compared to other cell lines (*P* < 0.05; Fig. 3D; Table S5). Lastly, the V55Y mutation inhibited propagation of all but one MSA sample (*P* = 0.26; Fig. 3E; Table S5), suggesting residue V55 plays a stronger role in forming the protofilament interface than G36 or V37 in MSA fibrils.

Finally, to disrupt the twister conformation with minimal impact on MSA replication, we mutated the valine at position 74 to either an isoleucine or a proline. Consistent with this objective, Maestro modeling indicated that both the V74I and V74P mutations would reduce the total energy of the α-synuclein conformation, though the effect of the V74P mutation was much stronger, suggesting MSA should be able to propagate (Table 2). However, *in vitro* studies showed that the V74I mutation blocks MSA replication (*P* = 0.0499; Fig. 3F; Table S5). Notably, while we detected a statistical difference between control and MSA samples in this cell line, the minimal effect size between the two groups indicates this statistical change is not biologically relevant. By comparison, the cell assay data showed that the V74P mutation merely blunts MSA replication (*P* < 0.05; Fig. 3G; Table S5) compared to the WT cells. This was unexpected given the substantial decrease in total energy predicted by Maestro.

It is worth noting that when we used Maestro to model the effect of mutations that were designed based on protein structures, it variably predicted our *in vitro* results. For example, while there was largely agreement between the analyses for the E61Q, V66F, and V55Y mutations, we saw less consistency between the predicted data and the biological effects of the G36K, V37F, V74I, and V74P mutations. One possible explanation for the G36K and V37F discrepancy is the fact that these two mutations are located at the end of subunit B, which decreases the reliability of Maestro calculations due to the increased flexibility arising from the smaller number of interactions. However, given that there are sample-to-sample differences in the effect of some mutations, it is possible that the presence of additional α-synuclein conformations in the MSA patient samples confounds our *in silico* analyses.

### The E46K mutation blocks MSA replication by disrupting the E46/K80 salt bridge

Several of the α-synuclein cryo-EM structures, including the MSA filaments, contain Greek key motifs that are stabilized by an intermolecular salt bridge that typically forms between residues E46 and K80 (reviewed in Ref^24^). In light of our previous findings that the E46K mutation blocks MSA replication,^11, 13^ we predicted that this effect occurs because the two lysines repel one another when the salt bridge is lost, destabilizing the misfolded conformation.^28^ To test this hypothesis, we generated four novel mutations designed to disrupt the E46/K80 salt bridge at position K80. In these mutations, we swapped the positively charged lysine for a negatively charged glutamic acid (K80E), predicting this would create a similar repellant interaction as the E46K mutation. We anticipated that the polar but uncharged asparagine (K80N) and glutamine (K80Q) would allow us to test the role of the positive charge on the lysine in salt bridge formation, given that the asparagine and glutamine have similar R groups to the lysine. And, finally, the bulky hydrophobic tryptophan (K80W) was selected to crowd out the packing of the R groups, preventing α-synuclein from adopting the Greek key motif. Using Maestro, we first confirmed that the E46K mutation is predicted to have a destabilizing effect on both subunits in the MSA fibril structure, with an increase in 70.2 and 51.6 kcal/mol, respectively (Table 3). We then tested the effect of the four K80 mutations using the same approach. While K80E, K80Q, and K80W all showed an increase in total energy, the K80N mutation showed a decrease in total energy for subunit A but an increase for subunit B.

**Table 3.**
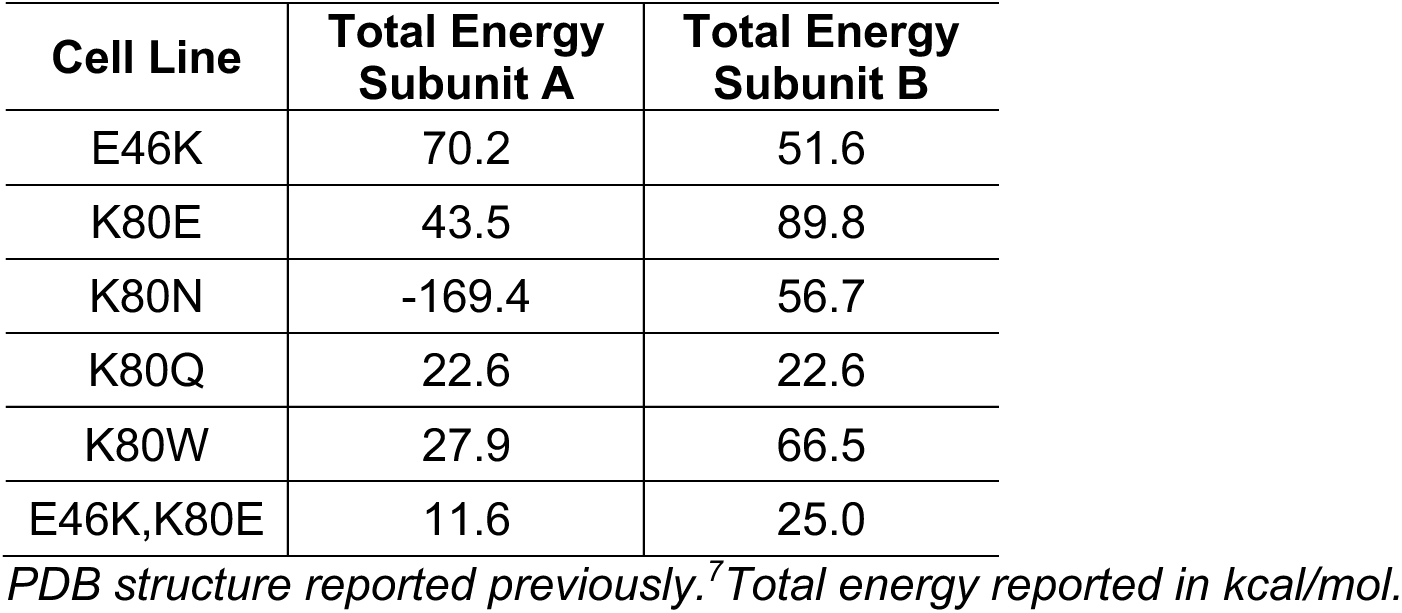
Total energy change after mutagenesis targeting the E46/K80 salt bridge in MSA structure 6XYQ.

Testing these mutations *in vitro*, we found that the K80E mutation strongly inhibits MSA replication (*P* = 0.49; Fig. 4A; Table S6), consistent with our E46K findings and our hypothesis that similar charges (in this case, a negative charge) would force apart the two stabilizing ends of the Greek key. By comparison, the K80N and K80Q mutations exert variable effects on MSA infection, with some MSA patient samples able to replicate using the mutant proteins while others were blocked (Figs. 4B & C; Table S6). The effect of the K80N mutation on MSA propagation was more consistent (*P* < 0.05), whereas the MSA samples showed a greater range in the ability to replicate using α-syn*K80Q (*P* = 0.30). Interestingly, the conflicted effect of the K80N mutation on the two subunits did not result in a completely inhibitory effect. These results suggest that the positive charge on the lysine is important for providing stability through the Greek key, but it is not always necessary for amplification. Additionally, we would expect the longer glutamine to occasionally replace the lysine in the MSA structures. However, the shorter asparagine would lack the length required to form other inter- and intramolecular interactions, leading to reduced stability compared to the K80Q mutation. Finally, the largest amino acid substitution, the K80W mutation, also largely blocked MSA prion replication (*P* = 0.06; Fig. 4D; Table S6). We hypothesize this occurs through steric hindrance created by the longer side chain with the large aromatic ring. These data are largely consistent with the predicted effects of each mutation by Maestro, underscoring the importance of the E46/K80 salt bridge in stabilizing the MSA fibril structure.

**Fig. 4.**
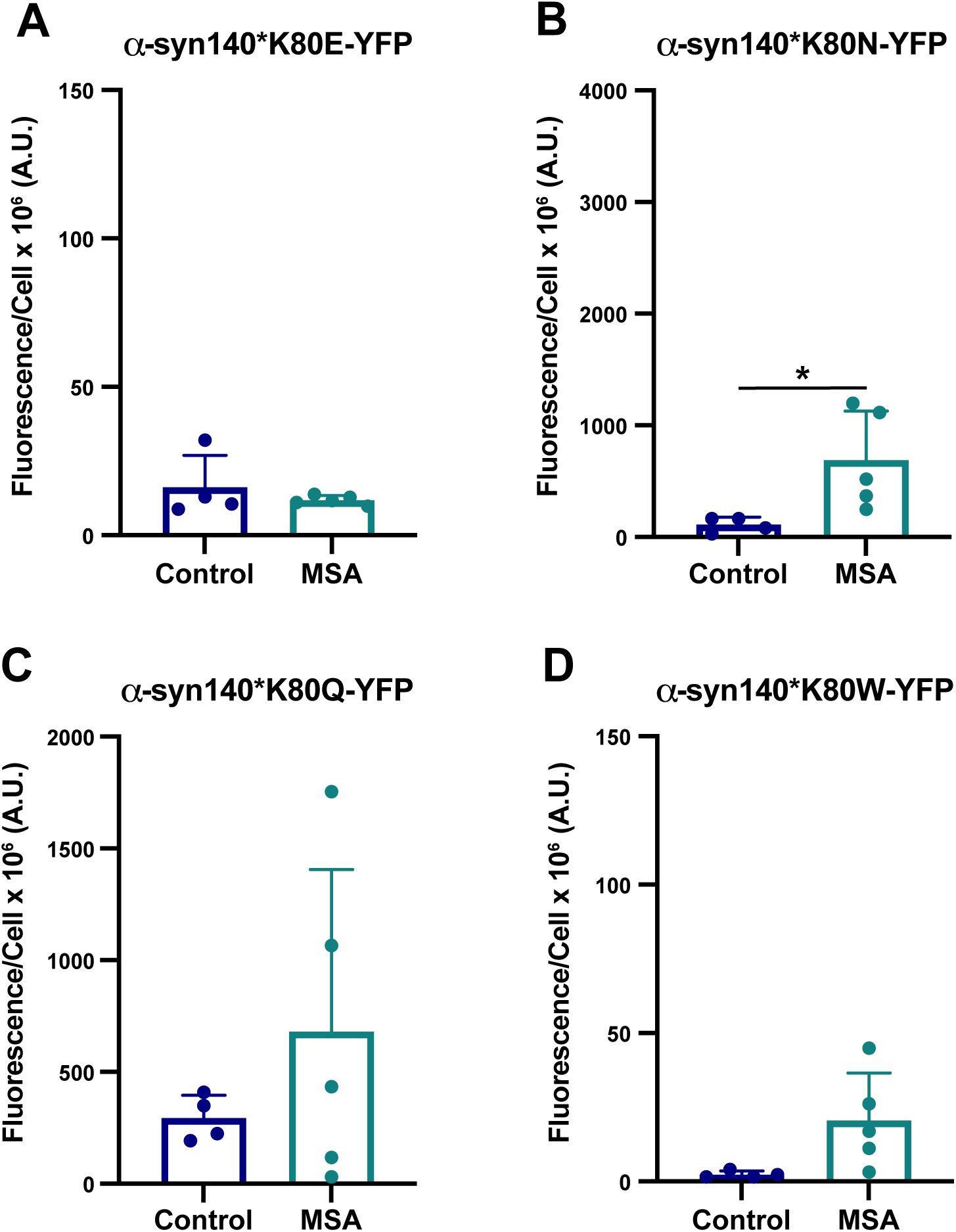
MSA prion replication requires the formation of the E46/K80 salt bridge. HEK293T cells expressing α-synuclein mutations at lysine 80 were used to determine the requirement of the E46/K80 salt bridge for MSA prion replication *in vitro*. Each cell line was infected with α-synuclein prions isolated from 4 control patient samples (C1, C2, C6, and C7) or 5 MSA patient samples (MSA2, MSA3, MSA6, MSA7, and MSA12) by phosphotungstic acid precipitation (× 10^6^ arbitrary units [A.U.]). (A) The K80E mutation (α-syn140*K80E-YFP) blocked MSA prion replication. (B & C) The K80N (α-syn140*K80N-YFP) and K80Q mutations (α-syn140*K80Q-YFP) exerted variable effects on MSA prion replication, with the K80N mutation having a stronger blunting effect overall than the K80Q mutation. (D) The K80W mutation (α-syn140*K80W-YFP) also prevented MSA prion replication. (* = *P* < 0.05)

Having established the importance of the E46/K80 salt bridge in MSA prion replication, we hypothesized that swapping the residues would enable salt bridge formation, thereby re-establishing MSA replication *in vitro*. To test this, we created a cell line expressing both the E46K & K80E mutations (E46K,K80E), which we then incubated with MSA patient samples. Unexpectedly, the E46K,K80E cell line also prevented MSA replication (*P* = 0.91; Fig. 5; Table S6), suggesting that the presence of the glutamic acid and lysine alone is not sufficient to enable MSA propagation. To determine what may be contributing to this result, we mutated both residues in the MSA fibril conformations in Maestro and found that the double mutation had a slight increase in total energy compared to the unmutated structure (Table 3). While this overall change is smaller than the individual mutations, the destabilizing effect in both side chains likely contributes to the *in vitro* measurements.

**Fig. 5.**
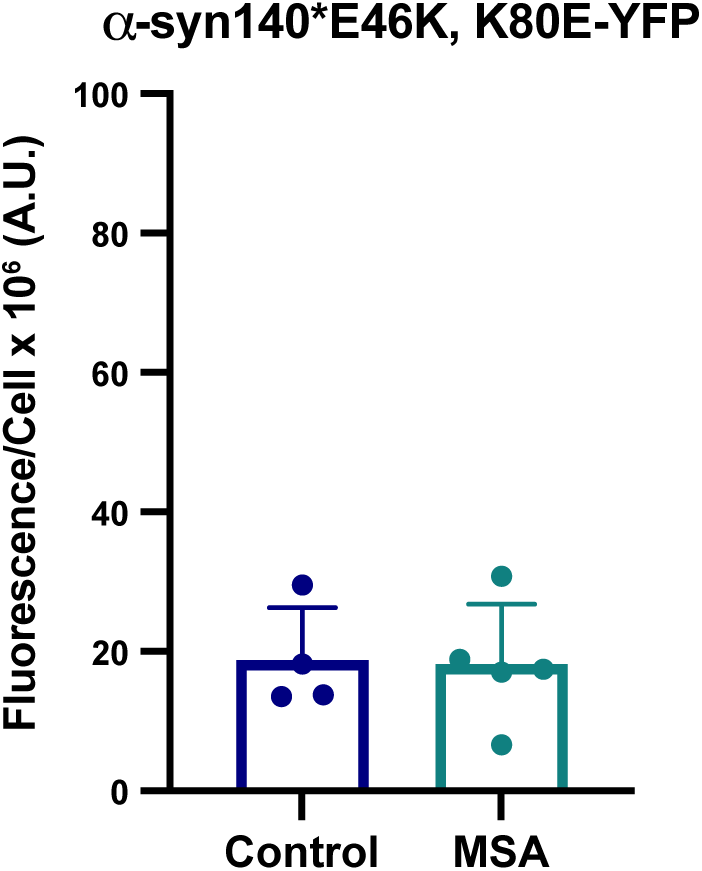
The E46K,K80E double mutation does not rescue MSA prion replication *in vitro*. HEK293T cells expressing the E46K & K80E double mutation in α-synuclein was used to determine if residue swapping can rescue MSA prion propagation. The cells were infected with α-synuclein prions isolated from 4 control patient samples (C1, C2, C6, and C7) or 5 MSA patient samples (MSA2, MSA3, MSA6, MSA7, and MSA12) by phosphotungstic acid precipitation (× 10^6^ arbitrary units [A.U.]). However, the MSA patient samples could not replicate in the α-syn140*E46K,K80E-YFP cells.

## DISCUSSION

Beginning with the solid-state NMR structure of α-synuclein fibrils reported by Tuttle *et al*., in 2016,^29^ the number of reported structures deposited in Protein Data Bank (PDB) has grown substantially due to advances in cryo-EM enabling the resolution of fibrillar structures made from recombinant protein^25–27, 30–37^ or isolated from human patient samples.^7, 8, 38^ However, as our understanding of the conformations that α-synuclein can adopt has increased, the resulting biological consequences of these structural differences have not been elucidated. Here, to start addressing this pivotal question for the α-synuclein conformations in MSA, we paired an *in silico* analysis of targeted mutagenesis with *in vitro* assays to experimentally validate the effect of each mutation on the ability of MSA prions to replicate. In addition to finding that our computational analyses using Maestro were typically able to predict the effect of a point mutation on α-synuclein misfolding, we established a panel of cell lines that can be used to define the biological properties of MSA prions. Moreover, the varied ability of each human patient sample to replicate using these cells suggests there may be multiple α-synuclein strains underlying disease, some of which appear to be biologically incompatible with the previously reported structures.^7^

Prior to the Goedert and Scheres labs determining the α-synuclein structures in MSA^7^ and LB diseases,^8^ we used our cell-based assay to investigate the ability of MSA prions to replicate *in vitro* using substrate containing the PD-causing mutations A30P, E46K, and A53T.^11^ The strong inhibitory effect of the E46K mutation on MSA propagation added to the growing biological and biochemical data indicating that the two diseases are caused by distinct α-synuclein strains,^10, 20, 21, 39^ but it was the structural determination that provided the most definitive evidence for this hypothesis. In light of these differences, we began using the protein structures as an opportunity to develop hypotheses about the effect of specific mutations on MSA replication (Fig. 1). Starting with the growing number of PD-causing mutations, we sought to further understand the biological differences between the MSA and Lewy strains by modeling the effect of the more recently discovered PD mutations on α-synuclein misfolding in MSA (Table 1 & Fig. 2). While our Maestro analyses indicated that the A30G, H50Q, and G51D mutations should be well tolerated (noting that we had limited data for A30G given its inconsistent presence in the templating region), we found that the A30G mutation largely inhibited MSA replication whereas it was mostly permissive in the presence of the H50Q and G51D mutations. It is important to note that both H50 and G51 are part of a heterodimeric pocket containing an unknown density, or co-factor. The Maestro minimizations were conducted on the published structures with no external molecules to balance the positive charge in this reason. As a result, we have low confidence in the H50Q and G51D predictions. By comparison, both the A53E and A53V mutations were predicted to be energetically unfavorable in subunit A, but had opposite effects on subunit B, with the A53V mutation reducing total energy. In line with the subunit B analyses, only the A53E mutation inhibited MSA propagation. Finally, the effect of the T72M mutation was more disfavored, which is consistent with the dampened *in vitro* replication observed.

We were particularly interested in evaluating the effect of the G51D and A53E mutations on MSA replication due to reports showing that patients with these mutations develop both LB and glial pathology, the latter of which resembles the GCIs in MSA patients.^16, 17^ Anticipating these mutations would exert a similar effect on MSA replication, we were surprised to find that the A53E mutation completely inhibited α-synuclein misfolding *in vitro* (Fig. 2G). Transmission studies using tissue from patients with the A53E mutation have not been reported, however, recent data from Lau *et al.* demonstrated that intracranial injection of patient samples carrying the G51D mutation can induce subclinical replication and pathological accumulation of α-synuclein in the brains of TgM83^+/-^ mice.^40^ The lack of observable neurological disease in these studies is consistent with previous efforts to transmit PD patient samples to the same mouse model.^20^ Lau *et al*. also found that the G51D patient samples behaved more like PD than MSA in an RT-QuIC assay optimized to detect either PD or MSA α-synuclein. These findings, combined with our cellular data, suggest that the glial co-pathology in the G51D and A53E patients is caused by a different α-synuclein strain than is present in MSA, and that the A53E mutation is incompatible with MSA replication.

In addition to investigating the effect of PD mutations on MSA propagation, we used the reported MSA and PFF structures to design non-pathogenic mutations with the goal of disrupting α-synuclein misfolding into specific conformations (Table 2 & Fig. 3). As expected, the E61Q and V66F mutations, which were designed to interfere with the rod and twister PFF conformations (residues noted in purple in Figs. 1C & D), largely facilitated MSA replication, as did the V74P mutation though with a blunted response. Unexpectedly, the V74I mutation blocked MSA propagation. This is particularly interesting given that our Maestro analyses suggested that both the V74I and V74P mutations would reduce the total energy of the structure. By comparison, the G36K, V37F, and V55Y mutations were all designed to interfere with the protofilament interface in the MSA structures, however, only the V55Y mutation inhibited propagation of most MSA patient samples. It should also be noted that residue V55 is located next to a small unfilled pocket in the cryo-EM structure, and it unclear if that pocket is filled with something exogenous. As a result, we were unable to account for these potential interactions in our calculations. While analyzing our cell assay data, we noticed that in some cell lines, such as V55Y, one of the MSA patient samples appeared as an outlier. We also noticed that there was often a large range in replication ability across the patient samples, as is seen in the V66F and G36K data. To determine if this variability is due to differences in titer, or the amount of replication competent α-synuclein present in each sample, we created a heat map showing cell infection relative to the average of the control samples for each patient sample on every cell line tested (Table 4). We observed that a single patient sample is neither responsible for the lowest nor the highest infection across the cell lines, suggesting that our findings are not due to differences in titer. Instead, these data suggest the potential presence of multiple α-synuclein strains in the MSA patient samples. Given that some mutations tested, for example V66F, should have no impact on MSA replication based on the location of the residue in the reported structures, and others, such as V55Y, should inhibit α-synuclein misfolding, it is highly likely that these unreported strains have conformations that are distinct from reported structures (both residues are highlighted in purple in Fig. 1A).

**Table 4.**
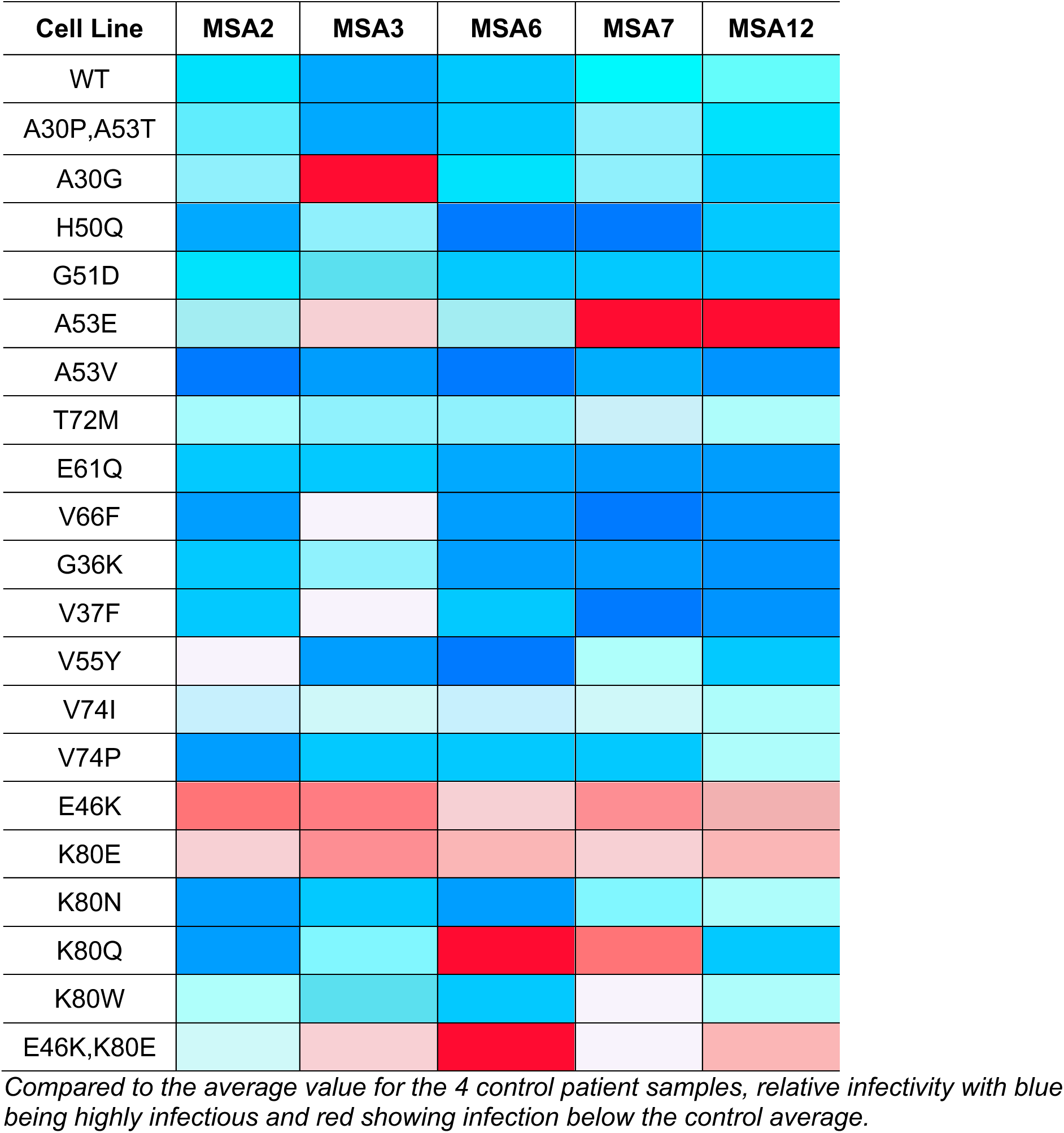
Heat map of MSA patient sample cell infection relative to control.

Our previous finding that the E46K mutation inhibits MSA replication is consistent with research on the prion protein (PrP), which has shown that a single residue change in the protein sequence can inhibit the conversion of cellular PrP^C^ into pathogenic PrP^Sc^. Notably, a single residue change is responsible for the species barrier creating resistance of the *Canidae* family to prion diseases,^41–43^ and for conferring resistance of sheep to scrapie^44, 45^ and humans to Creutzfeldt-Jakob disease.^46^ At the time we discovered the effect of the E46K mutation on MSA replication, we relied on the solid-state NMR structure of α-synuclein PFFs^29^ to interpret our findings. In this structure, a Greek key motif is stabilized by a salt bridge between residues E46 and K80, leading to our hypothesis that the E46K mutation prevents salt bridge formation.^47^ The more recent resolution of α-synuclein fibrils isolated from MSA patient samples confirmed that this salt bridge stabilizes the MSA conformations,^7^ supporting our initial hypothesis. To test this, we designed four mutations targeting the second half of the salt bridge (residue K80) and found that the K80E and K80W mutations blocked MSA propagation (Fig. 4). With the goal of re-establishing the salt bridge to rescue MSA replication, we then built cells expressing the double E46K,K80E mutations, and were surprised to find that MSA transmission remained inhibited (Fig. 5). Modeling the effect of this residue swap using Maestro showed that the double mutation results in a small increase in total energy for both subunits due to an angular salt bridge with minimal orbital overlap, likely resulting in the inhibitory effect observed *in vitro*. Notably, the E46/K80 salt bridge is an intermolecular interaction, rather than intramolecular, meaning that the salt bridge extends between two layers of the protofilament. Given the strong effect of the E46K and K80E mutations on blocking MSA propagation, this salt bridge may act as a key area on the fibril surface for driving self-templating, serving as an anchoring point to facilitate α-synuclein conversion.

An important finding from this work is that protein modeling software is not always capable of accurately predicting the effect of each mutation on MSA strain biology. For example, as was the case for residues G36, V37, H50, and G51, the location of the residue within the templating region contributes to the reliability of the modeling data. There are three important caveats to our approach that may contribute to these discrepancies. First, our methods used implicit waters in the force field calculations instead of explicit waters. Second, there are pockets in the MSA fibril structures, some of which are known to be occupied by non-protein densities, however, because it is unknown what molecules are present in these pockets, we are unable to account for their interactions with mutated residues. And third, we are using thermodynamic calculations to try to predict a kinetic process. While Maestro can be used to assess the capacity of an established structure to house a mutated residue, it is unable to predict the ability of the mutant substrate to misfold into a specific conformation.

Attempting to address this limitation, our initial approach to modeling the effect of each mutation using Maestro focused on mimicking the experimental design of our *in vitro* studies by only mutating 3 layers of the protein fibril. The goal was to computationally recapitulate the effect of self-templating on the catalytic surface of the MSA fibril using the mutant α-synuclein expressed in each cell line. However, we quickly discovered that this approach failed to fully capture the consequences of each mutation on the change in total energy, making it difficult to assess the likely effect of each mutation on MSA replication *in vitro*. Instead, when we expanded our analyses to mutate 6 layers of the protofilament, the effect of each mutation was much more apparent, increasing the applicability of our *in silico* modeling to interpreting our *in vitro* data (Table 5). The problem with mutating just 3 layers of the protein fibril may, in part, be an unintended consequence of our control mutation (often E83X, which was chosen to minimize interactions with neighboring residues). This step was required to keep the amino acids identical for the purpose of comparing total energies. Intramolecular interactions between the sidechains of WT and mutant amino acids were often energetically favorable. The control mutation doubles the energy of these interactions, distorting the energy calculations given that the control mutations are not present in the cell models.

**Table 5.**
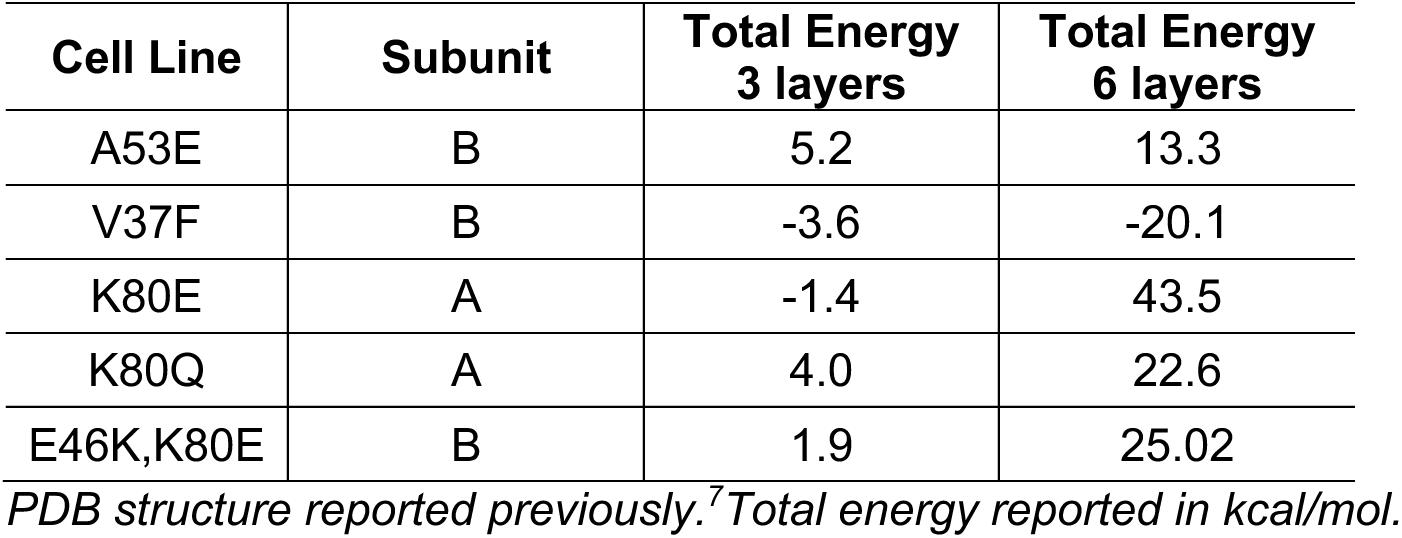
Total energy change after mutagenesis of 3 versus 6 protein layers in MSA structure 6XYQ.

A final critical limitation to our *in silico* approach is the inability to predict deformed templating of a prion strain. In cases where Maestro predicted a mutation would increase total energy but templating was still permitted *in vitro*, such as the V37F mutation, it is possible that a slight alteration in the templating surface occurs to facilitate incorporation of the mutant residue. Deformed templating is thought to be a contributor to prion strain adaptation when strains are successfully transmitted to hosts with a different genotype or across species barriers.^48–51^ In these cases, future studies should investigate maintenance of strain properties after passaging to determine if deformed templating occurred, and if so, how the altered protein structure impacts prion biology.

The remarkable advances made in structural biology methodology, particularly for prionogenic proteins, have had a tremendous impact on our understanding of protein misfolding and strain biology. The availability of these structures now serves as an opportunity for hypothesis development focused on elucidating the structure-function relationship underlying the ability of each prion strain to exert unique biological consequences in disease. However, the nature of structural techniques limits our understanding of fibril formation to a single snapshot from the brain of a deceased patient, failing to capture the dynamic processes that lead to the final fibrillar product. In this study, we paired structural analysis with *in vitro* models of α-synuclein misfolding to directly test the effect of targeted mutations on MSA propagation with the goal of understanding how α-synuclein misfolds into distinct conformations. Having shown that MSA prion replication is incompatible with some of the familial PD *SNCA* mutations, we demonstrated that the mechanism underlying the E46K-driven inhibition is due to disruption of a critical salt bridge between residues E46 and K80. Moreover, using novel mutations designed to perturb protein misfolding, we find evidence suggesting additional unresolved α-synuclein strains are present in MSA patient samples. In addition to furthering our understanding of α-synuclein misfolding in MSA, this work has established a powerful panel of α-syn140-YFP cell lines that can be used to interrogate MSA strain biology.

## ACKNOWLEDGMENTS

The authors thank Scott Horowitz for his assistance with the computational modeling.

