## Supplemental for "Structurally targeted mutagenesis identifies key residues supporting α-synuclein misfolding in multiple system atrophy"

### Journal of Parkinson's Disease

Patricia M. Reis[1,2], Sara A. M. Holec[1,3#], Chimere Ezeiruaku[1#†], Matthew P. Frost[1#‡], Christine K. Brown[1ˆ], Samantha L. Liu[1ƒ], Steven H. Olson[4], and Amanda L. Woerman[1,3*]

[1]Department of Biology and Institute for Applied Life Sciences, University of Massachusetts Amherst, Amherst, MA, USA; [2]Neuroscience and Behavior Graduate Program, University of Massachusetts Amherst, Amherst, MA, USA; [31]Department of Microbiology, Immunology, and Pathology, Prion Research Center, Colorado State University, Fort Collins, CO, USA; [4]Conrad Prebys Center for Chemical Genomics, Sanford Burnham Prebys Medical Discovery Institute, San Diego, CA, USA.

[#]Authors contributed equally to the manuscript.

[†]Current affiliation: Department of Surgery, Division of Abdominal Transplant Surgery, Stanford University School of Medicine, Palo Alto, CA, USA.

[‡]Current affiliation: Neuroscience Department, UConn Health, Farmington, CT, USA.

[ˆ]Current affiliation: Department of Biomedical Engineering, University of Massachusetts Amherst, Amherst, MA, USA.

[ƒ]Current affiliation: Department of Biochemistry & Cell Biology, Dartmouth College, Hanover, NH, USA.

*Corresponding author: Amanda L. Woerman, PhD, Department of Department of Microbiology, Immunology, and Pathology and Prion Research Center, Colorado State University, 300 West Lake St, Fort Collins, CO 80526,.

### MATERIALS AND METHODS

#### Human patient neuropathology

Neuropathology assessment by the Massachusetts Alzheimer's Disease Research Center (MADRC) Brain Bank was performed using human tissue samples fixed in 10% (vol/vol) neutral buffered formalin and coronally sectioned. The fixed tissue was evaluated histologically using a set of blocked regions representative of a variety of neurodegenerative diseases. All blocks were stained with H&E and luxol fast blue. Selected blocks were used for immunohistochemical staining for α-synuclein, β-amyloid, and phosphorylated tau. A confirmed MSA diagnosis required the presence of glial cytoplasmic inclusions.[1]

Human samples provided by the Sydney Brain Bank were fixed in 15% (vol/vol) buffered formalin [39% (vol/vol) aqueous formaldehyde solution] for 2 weeks before sectioning. Standard neuropathological assessment was done using H&E-stained sections, and a modified Bielschowsky silver stain was used to identify Alzheimer-type pathologies. Immunostaining for phosphorylated α-synuclein (BD Biosciences USA; 1:7,000), phosphorylated tau (AT8, Thermo Scientific USA; 1:1,000), and β-amyloid (Dako Denmark; 1,500) was performed.

**Table S1. Patient sample information.**

| Patient | Disease | Age at Death | Sex | Brain Region | Brain Bank |
|---|---|---|---|---|---|
| C1 | CAA | 84 | F | Midbrain | MADRC[a] |
| C2 | None | 90+ | M | Midbrain | MADRC[a] |
| C6 | None | 79 | M | Pons | NIH NeuroBioBank |
| C7 | None | 70 | F | Pons | NIH NeuroBioBank |
| MSA2 | MSA | 68 | M | Midbrain | MADRC[a] |
| MSA3 | MSA | 61 | M | Pons | Sydney Brain Bank |
| MSA6 | MSA | 71 | F | Pons | Sydney Brain Bank |
| MSA7 | MSA | 72 | F | Pons | Sydney Brain Bank |
| MSA12 | MSA | 77 | F | Midbrain | MADRC[a] |

[a] *Massachusetts Alzheimer's Disease Research Center*

**Table S2. Cell assay experimental conditions.**

| Cell Line | Cells per well | Lipofectamine (%) | Sample dilution in DPBS |
|---|---|---|---|
| WT | 4000 | 1.5 | 1:5 |
| A30P,A53T | 3500 | 1.5 | 1:10 |
| A30G | 2500 | 1 | 1:5 |
| H50Q | 3250 | 2.5 | 1:5 |
| G51D | 3500 | 2 | 1:4 |
| A53E | 3500 | 1.5 | 1:10 |
| A53V | 3250 | 1.5 | 1:7 |
| T72M | 4000 | 2 | 1:10 |
| E61Q | 3500 | 2 | 1:10 |
| V66F | 4250 | 1 | 1:10 |
| G36K | 3000 | 1.5 | 1:10 |
| V37F | 4000 | 1 | 1:20 |
| V55Y | 3500 | 1 | 1:20 |
| V74I | 2750 | 1 | 1:20 |
| V74P | 4500 | 1.5 | 1:10 |
| E46K | 3500 | 1 | 1:10 |
| K80E | 3250 | 1 | 1:20 |
| K80N | 3500 | 1 | 1:10 |
| K80Q | 3500 | 1.5 | 1:10 |
| K80W | 3250 | 1 | 1:10 |
| E46K, K80E | 3500 | 1 | 1:20 |

**Table S3. Mutations used to control for changes in total energy in Maestro calculations.**

| Cell Line | Correcting Mutation |
|---|---|
| A30G | A85G |
| H50Q | E83Q |
| G51D | E83D |
| A53E* | E83A |
| A53V | E83V |
| T72M | E83M |
| E61Q | E83Q |
| V66F | E83F |
| G36K | E83K |
| V37F | E83F |
| V55Y | E83Y |
| V74I | E83I |
| V74P | V16P |
| E46K | K34A |
| K80E | E83K |
| K80N | E83N |
| K80Q | E83Q |
| K80W | E83W |
| E46K, K80E | None |

**The absence of the negatively charged non-protein density in the PDB structure freed K43 to make a salt bridge with A53E. To prevent this, the K43A mutation was inserted in both the control and mutant calculations.*

**Table S4. Cell infection data in α-syn140-YFP cells expressing PD-causing mutations.**

| Cell Line | C1 | C2 | C6 | C7 | MSA2 | MSA3 | MSA6 | MSA7 | MSA12 |
|---|---|---|---|---|---|---|---|---|---|
| WT | 120 ± 74 | 320 ± 140 | 360 ± 180 | 110 ± 37 | 2800 ± 1000 | 4200 ± 860 | 3200 ± 700 | 1500 ± 690 | 1400 ± 450 |
| E46K | 0.2 ± 0.2 | 1.1 ± 1.1 | 1.7 ± 0.7 | 0.7 ± 0.8 | 6.8 ± 5.0 | 11 ± 2.1 | 38 ± 20 | 15 ± 4.8 | 24 ± 15 |
| A30P,A53T | 16 ± 9.6 | 11 ± 5.2 | 13 ± 6.6 | 21 ± 8.2 | 1400 ± 430 | 3700 ± 550 | 3100 ± 660 | 610 ± 300 | 1600 ± 550 |
| A30G | 18 ± 10 | 5.9 ± 6.1 | 3.7 ± 3.8 | 3.9 ± 3.5 | 15 ± 9.0 | 4.3 ± 3.2 | 37 ± 19 | 15 ± 12 | 110 ± 25 |
| H50Q | 46 ± 33 | 230 ± 120 | 150 ± 68 | 120 ± 81 | 4500 ± 1700 | 440 ± 70 | 4900 ± 2100 | 5100 ± 1500 | 2500 ± 850 |
| G51D | 14 ± 10 | 6.8 ± 6.2 | 19 ± 16 | 22 ± 7.1 | 210 ± 110 | 92 ± 46 | 470 ± 250 | 380 ± 120 | 330 ± 280 |
| A53E | 23 ± 20 | 97 ± 79 | 16 ± 10 | 18 ± 13 | 98 ± 54 | 45 ± 25 | 100 ± 58 | 18 ± 11 | 23 ± 18 |
| A53V | 520 ± 520 | 430 ± 190 | 530 ± 290 | 190 ± 39 | 12000 ± 3200 | 7000 ± 2600 | 15000 ± 2700 | 5800 ± 1900 | 8000 ± 2100 |
| T72M | 33 ± 16 | 13 ± 10 | 11 ± 10 | 22 ± 15 | 86 ± 54 | 100 ± 61 | 90 ± 51 | 28 ± 14 | 55 ± 24 |

*Mean fluorescence/cell ± standard deviation (× $10^6$ arbitrary units [A.U.]).*

**Table S5. Cell infection data in α-syn140-YFP cells expressing structurally informed mutations.**

| Cell Line | C1 | C2 | C6 | C7 | MSA2 | MSA3 | MSA6 | MSA7 | MSA12 |
|---|---|---|---|---|---|---|---|---|---|
| E61Q | 37 ± 29 | 30 ± 4.2 | 37 ± 29 | 19 ± 7.9 | 400 ± 90 | 450 ± 120 | 700 ± 190 | 1200 ± 480 | 990 ± 110 |
| V66F | 160 ± 210 | 1200 ± 600 | 35 ± 42 | 710 ± 1400 | 2000 ± 1200 | 500 ± 370 | 2100 ± 1200 | 7300 ± 5800 | 4700 ± 1900 |
| G36K | 20 ± 6.8 | 22 ± 7.6 | 41 ± 12 | 26 ± 9.5 | 240 ± 74 | 110 ± 38 | 430 ±140 | 650 ± 180 | 1200 ± 490 |
| V37F | 160 ± 210 | 1200 ± 600 | 35 ± 42 | 710 ± 1400 | 2000 ± 1200 | 500 ± 370 | 2100 ± 1200 | 7300 ± 5800 | 4700 ± 1900 |
| V55Y | 93 ± 62 | 34 ± 34 | 49 ± 18 | 39 ± 16 | 55 ± 30 | 2100 ± 730 | 12000 ± 3100 | 260 ± 140 | 730 ± 200 |
| V74I | 1.8 ± 1.7 | 1.7 ± 2.5 | 0.9 ± 1.5 | 2.4 ± 3.6 | 7.1 ± 5.6 | 4.9 ± 2.0 | 6.5 ± 7.1 | 5.2 ± 3.3 | 18 ± 9.2 |
| V74P | 9.4 ± 7.8 | 12 ± 6.8 | 6.8 ± 5.8 | 11 ± 4.0 | 120 ± 39 | 68 ± 18 | 49 ± 13 | 51 ± 27 | 38 ± 16 |

*Mean fluorescence/cell ± standard deviation (× $10^6$ arbitrary units [A.U.]).*

**Table S6. Cell infection data in α-syn140-YFP cells expressing mutations targeting the E46/K80 salt bridge.**

| Cell Line | C1 | C2 | C6 | C7 | MSA2 | MSA3 | MSA6 | MSA7 | MSA12 |
|---|---|---|---|---|---|---|---|---|---|
| K80E | 32 ± 34 | 8.8 ± 11 | 11 ± 8.7 | 13 ± 12 | 13 ± 14 | 9.8 ± 8.7 | 11 ± 5.3 | 14 ± 12 | 12 ± 8.6 |
| K80N | 29 ± 26 | 170 ± 77 | 170 ± 110 | 81 ± 37 | 1100 ± 470 | 520 ± 210 | 1200 ± 450 | 370 ± 230 | 250 ± 33 |
| K80Q | 190 ± 120 | 350 ± 210 | 220 ± 180 | 410 ± 250 | 1800 ± 2300 | 430 ± 200 | 31 ± 29 | 120 ± 46 | 1100 ± 220 |
| K80W | 2.3 ± 1.4 | 1.5 ± 1.5 | 4.0 ± 1.8 | 1.5 ± 0.8 | 17 ± 4.7 | 26 ± 11 | 45 ± 22 | 3.2 ± 2.2 | 11 ± 4.6 |
| E46K,K80E | 14 ± 6.1 | 14 ± 6.4 | 30 ± 12 | 18 ± 13 | 31 ± 26 | 18 ± 12 | 6.7 ± 2.3 | 19 ± 6.8 | 17 ± 6.8 |

*Mean fluorescence/cell ± standard deviation (× $10^6$ arbitrary units [A.U.]).*
